# Expanding an expanded genome: long-read sequencing of *Trypanosoma cruzi*

**DOI:** 10.1101/279174

**Authors:** Luisa Berná, Matías Rodríguez, María Laura Chiribao, Adriana Parodi-Talice, Sebastián Pita, Gastón Rijo, Fernando Alvarez-Valin, Carlos Robello

## Abstract

Although the genome of *Trypanosoma cruzi*, the causative agent of Chagas disease, was first made available in 2005, with additional strains reported later, the intrinsic genome complexity of this parasite (abundance of repetitive sequences and genes organized in tandem) has traditionally hindered high-quality genome assembly and annotation. This also limits diverse types of analyses that require high degree of precision. Long reads generated by third-generation sequencing technologies are particularly suitable to address the challenges associated with *T. cruzi´s* genome since they permit directly determining the full sequence of large clusters of repetitive sequences without collapsing them. This, in turn, allows not only accurate estimation of gene copy numbers but also circumvents assembly fragmentation. Here, we present the analysis of the genome sequences of two *T. cruzi* clones: the hybrid TCC (DTU TcVI) and the non-hybrid Dm28c (DTU TcI), determined by PacBio SMRT technology. The improved assemblies herein obtained permitted us to accurately estimate gene copy numbers, abundance and distribution of repetitive sequences (including satellites and retroelements). We found that the genome of *T. cruzi* is composed of a "core compartment" and a "disruptive compartment" which exhibit opposite gene and GC content composition. New tandem and disperse repetitive sequences were identified, including some located inside coding sequences. Additionally, homologous chromosomes were separately assembled, allowing us to retrieve haplotypes as separate contigs instead of a unique mosaic sequence. Finally, manual annotation of surface multigene families MUC and trans-sialidases allows now a better overview of these complex groups of genes.

## Introduction

The year 2005 represents a landmark in the study of trypanosomatid biology with the simultaneous publication of the *Leishmania major, Trypanosoma brucei and Trypanosoma cruzi* genomes; three species considered to be representative of the parasitic diversity of the group (1–3). The new information unraveled by these genomes opened a new era that allowed researches to conduct studies with unprecedented comprehensiveness. Several new types of analyses, such as large scale comparative genomics aimed at understanding common evolutionary basis of parasitism and pathogenesis, or searching for new vaccine candidates and drug targets (4,5) were made possible. These three genomes, however, were published with very different finishing degrees. The genome of *L. major* was precisely assembled and annotated. The *T. brucei* genome was also of very good quality but was mostly focused on megabase chromosomes, whereas minichromosomes were not included. Conversely, the assembly of the *T. cruzi* genome -CL Brener strain-was extremely fragmented (4098 contigs) with very few contigs exceeding 100 kb in size. In fact, only 12 contigs met this threshold and none was larger than 150 kb (3). Despite this degree of fragmentation, this draft genome was highly valuable because it led to the identification of several novel specie-specific multigene families and gave, for the first time, a draft overview of the chromosome architecture in *T. cruzi*. This draft, also made it possible to identify the vast majority of trypanosomatid conserved genes, as indicated by the fact that very few genes conserved between *Leishmania* and *T. brucei* were not found in the draft *T. cruz*i genome (4). Most likely this can be attributed to the fact that genes are short in trypanosomatids (since they lack introns), hence even moderately small contigs can contain complete coding sequences (CDS). The genome sequences of additional *T. cruzi* strains Dm28c (6), Sylvio X10/1 (7), and *T*. *c*. *marinkellei* (8) have been reported since this initial publication using new sequencing technologies such as Roche 454 alone or in combinations with other methodologies (Illumina and Sanger). A common feature of all the assemblies reported for these strains is again high fragmentation, which was even higher than that originally reported for CL Brener.

To tackle the problem of assembly fragmentation, Weatherly, Boehlke and Tarleton (9) used complementary source of data to scaffold *T. cruzi* (CL Brener strain) contigs aiming to recover full length chromosome sequences. Their approach used a combined strategy based on sequencing BAC ends, as well as their co-location on chromosomes, and synteny conservation with *T. brucei* and *Leishmania* chromosomes. This strategy enabled these authors not only to obtain an assembly that represented a substantial improvement in comparison to previous versions of the genome, but also to reconstruct chromosomes (with few gaps), with only a relatively minor portion of the genome remaining un-assigned to chromosomes. Nonetheless, the issue of assembly fragmentation is a limitation that poses a number of complications for diverse types of analyses that require high precision.

Third-generation sequencing technologies are particularly suitable to address the challenges associated with the peculiarities of the *T. cruzi’*s genome since they allow to obtain sequencing reads of more than 15 kb in average length and many much longer. This opens the possibility of directly determining the full sequence of large clusters of repetitive sequences (without collapsing them), as well as determining the single copy sequences that surround both sides of these clusters. Moreover, the collapse of tandemly arrayed gene copies erases any variability that these copies might eventually exhibit, thus precluding any assessment in which this variability may be relevant (for instance intra-cluster differential gene expression). As a consequence, assembly fragmentation is largely overcome. In fact, using this technology has made it possible to obtain much better quality genome assemblies (compared to pre-existing ones) in other parasitic protozoa that also bear a highly repetitive genome, such as *Plasmodium*, (10). More recently the full length chromosome sequences (without gaps and collapsed segments) were obtained in *P. malariae* and *P. ovale* (11). Furthermore, this technology allows to obtain separated assemblies of homologous chromosomes, such that haplotypes can be retrieved as separated contigs/scaffolds instead of a unique mosaic sequence. This feature is particularly desirable in genomes with high level of heterozygocis, as in the case of some *T. cruzi* hybrid strains that have arisen from relatively divergent ancestors, in which the divergence between haplotypes can be as high as 5%. Needless to say that a single mosaic assembly, representing a mixture of two parental sequences, can produce distortions in several types of analyses such as detecting recombination or aneuploidies.

In order to improve the genome information of this parasite we report here the genome sequence and assembly of two *T. cruzi* strains,TCC and Dm28c using long read sequencing. Our results not only significantly improve the quality of the genomic sequence and annotation available for this parasite, but also reveal multiple unique genomic features previously underestimated, such as gene copy numbers, plausible pathways of evolution of hybrid strains, whole genome compartmentalization into regions of specific sequence composition, and biased distribution of genes. Our results bring us one step closer to fully understanding *T. cruzi´s* biology by opening new avenues to evolutionary and population structure studies, as well as, allowing more precise gene number determination, and consequently, more efficient genetic manipulation.

## Results and discussion

### Genome assembly

To obtain a more complete assembly of the complex *T. cruzi* genome using the PacBio technology library preparation included fragmentation and size selection with a cutoff of 10Kb. This size threshold also prevents the inclusion of minicircles, which presence leads to a substantial reduction in sequencing depth of nuclear genomic DNA (they represent about 20% of total DNA). To include both clinically and evolutionarily relevant strains, we chose to sequence the hybrid TCC strain (Tc VI), derived from Tulahuen 2 and closely related to CL Brener (13), and the non-hybrid strain Dm28c (Tc I) (12).It should be mentioned that TcI is the most abundant DTU and has a wide distribution in the Americas, remaining as a pure line evolving separately from a common ancestor (Zingales et al Infect Genet Evol 2012) while TcVI, present in several countries of South America, is an hybrid of the two distinct lineages TcII and TcIII (41)

After filtering by read quality, we obtained 5.2 and 4.0 Gb of sequences comprising 343,384 and 261,398 reads for TCC and Dm28c, respectively (Table 1). The average read length was 16 Kb for both strains, and the longest reads were larger than 60Kb (Fig. 1). The availability of such long reads is essential to disentangle the sequence repetitions; a hallmark of *T. cruzi*. HGAPv.3 assembler (42) was used to correct and assemble the initial reads. *IPA* (15) was used to collapse overlapping contigs and cleanup the assembly incorporating illumina reads. The length of the reads enabled us to directly obtain the full sequence of large clusters of repetitive sequences without collapsing them, as well as to determine the non-repetitive sequences flanking those repeats. Illustrative examples of how PacBio reads allowed us to overcome some of the common complications associated with *T. cruzi* assemblies fragmentation and collapse of repeats-are presented in Fig. 2. The figure shows the comparison between chromosome 30-P from CL Brener (assembly reported in (9)) with TCC and Dm28C contigs reported here using long reads. In order to build the virtual chromosome 30-P (from CL Brenner) Weatherly et al. connected many contigs by segments of unknown sequence (filled with N letters, shadowed in green in Fig. 2(a)). These “unknowns” are fully resolved in our assemblies, both in TCC and Dm28c (Fig. 2(a)).

**Table 1.**
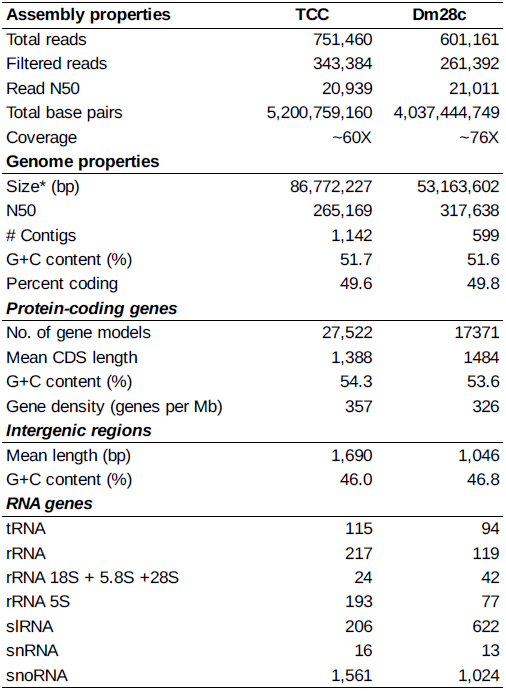
Summary of the assembly and annotation T. cruzi TCC and T. cruzi Dm28c genomes

**Fig. 1.**
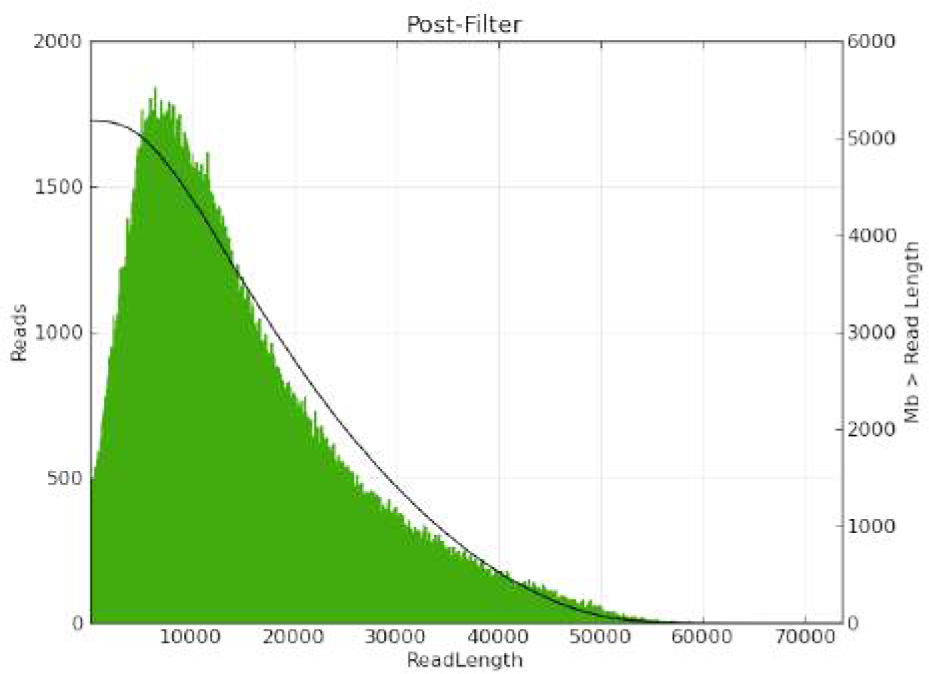
Length distribution of filtered reads from TCC strain.

**Fig. 2.**
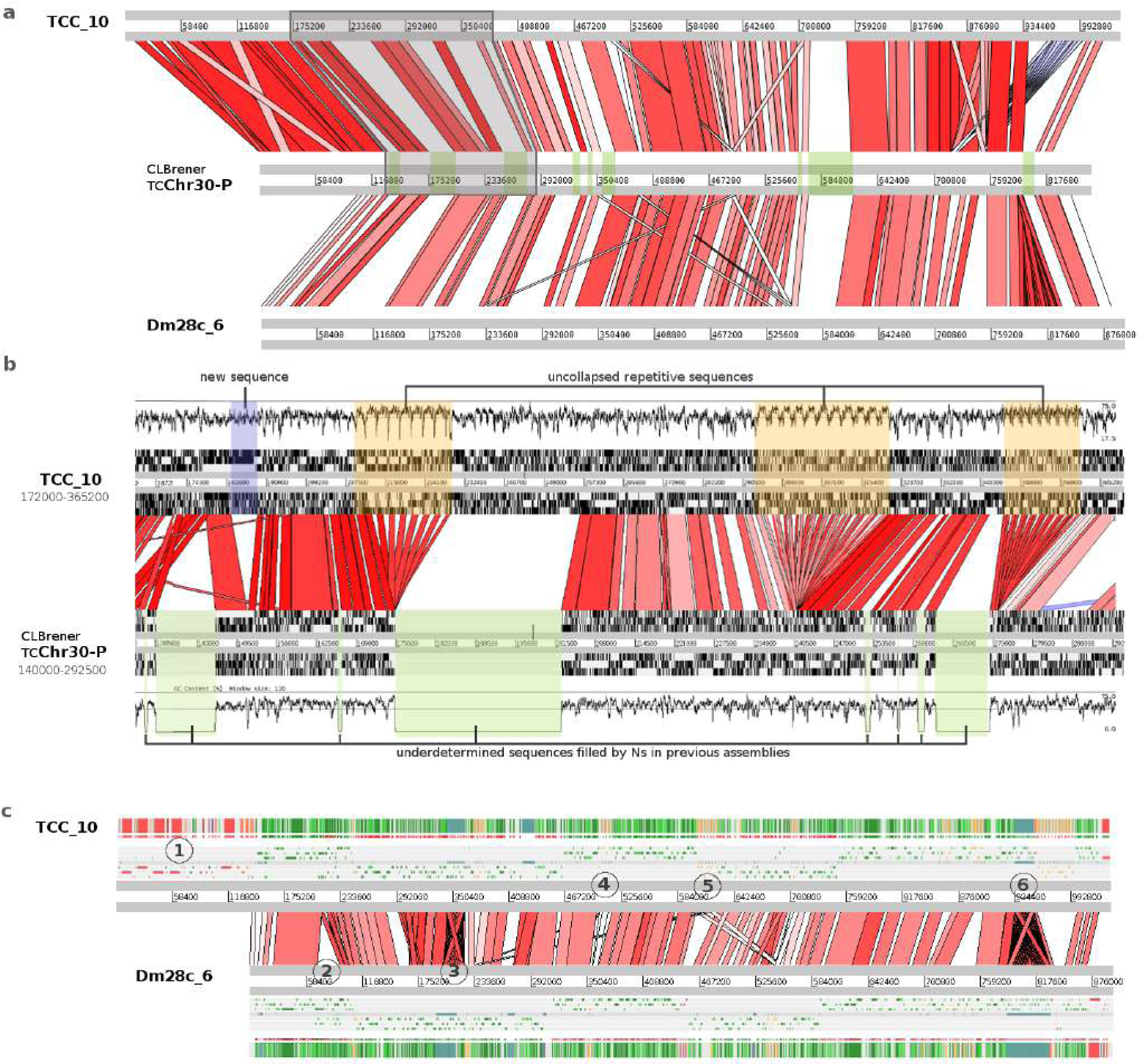
Chromosomal assembly improvements. **A)** ACT alignment of homologous chromosomes from three strains: TCC (contig TCC_10), Dm28c (contig Dm28c_6) and CL Brener (chromosome TcChr30-P). Previously undetermined sequences filled by Ns in CL Brener are marked in green. **B)** Magnification of a fragment of **A** (boxed and shadowed in gray). The six frames and the GC content of each chromosome are plotted. Previously collapsed repetitive sequences (boxed in orange) are disaggregated in the new assembly. **C)** Visualization of the alignment of the same homologous chromosome showing additional details in TCC and Dm28c. The color patterns in the annotation bars (bottom and top-most horizontal stripped bars) correspond to the annotation as they appear in the web interface (DGF1 in red, GP63 orange, RHS brown, conserved genes in green). The six reading frames are also shown. **1-** Terminal DGF-1 gene cluster present only in TCC. **2-** Non-homologous region present only in Dm28c. **3-** Repetitive region is present in both strains. **4-** Expansion of a GP63 cluster in TCC (four copies versus two copies in Dm28c). **5-** Strain specific amplifications of two different genes. There are seven GP63 copies (orange strips on the top annotation bar) in TCC but only one in Dm28c, moreover Dm28c contains four RHS copies in the same region. **6-** Repetitive element present in both genomes having fewer copies in TCC (20 copies in TCC and 44 copies in Dm28c). The segment is followed by another strains specific amplification consisting of a cluster of 14 GP63 genes in TCC and only one copy in Dm28c.

A second noticeable improvement is related to the repetitive sequences clusters, which were collapsed in the previous assembly, and now are disaggregated into the actual copy number (Fig. 2).This is more clearly shown in panel B from the same figure that shows a magnification of a fragment from the same CL Brener chromosome, containing several clusters of tandem repeats. The figure also shows that some of these collapsed clusters contributed to assembly fragmentation, since they are located in contig boundaries.

A comparison between the same chromosome of TCC and Dm28C (Fig. 2(c)) revealed additional aspects concerning cluster repeats and structural variation between strains. Specifically the figure shows copy number variations between the two strains in four groups (indicated by numbers in Fig. 2(c)) as well as strains specific insertions/deletions. Among the latter, it is worth mentioning a large segment located on the (left) telomere present in TCC and CL Brener, but absent in Dm28c, containing tandem repeats of some species specific genes (such as DGF1).

Overall, in terms of integrity (*i.e.* low fragmentation levels) the assemblies obtained for both strains, represent notorious improvements compared to previous ones (Table 1). Their N50 values were 265 and 317 Kb for TCC and Dm28c respectively, representing more than a ten-fold increase in this index. Furthermore, several contigs correspond to entire chromosomes (being the largest ones 1.65 Mb for Dm28c and 1.35 Mb for TCC). Other contigs are considerably smaller (50 kb) and are composed uniquely of a well known satellite sequence of 195 bp that encompasses more than 5% of the genome (see below). This extremely abundant repeat is one of the factors that contributes the most to assembly fragmentation.

Besides fragmentation, another important element to consider when evaluating assembly quality is its size. In the case of the Dm28c, the assembly size was 53.2 Mb, consistent with its haploid size as previously estimated using a fluorescent nucleic acid dye (43). Previous efforts to sequence Dm28c and Sylvio X10/1 strains (two closely related strains from TcI) using other technologies (6,7) resulted not only in assembly fragmentation but also in genome sizes of roughly one half of the actual haploid size (27Mb for Dm28c and 23Mb for Sylvio X10/1). This size underestimation is in all likelihood due to the limitations of short reads technologies for assembling complex genomes. Keeping this limitation in mind, Franzen *et al.* (2011) recalibrated their genome size estimation for Sylvio X10/1 strain to 44 Mb, extrapolating on the basis of non-assembled reads.

The TTC strain is closely related to CL Brener, hence, we expected their genomes to be very similar in terms of size and sequence composition. Indeed, sequence identity is higher than 99.7% over segments longer than 10 kb (Table S1). The CL Brener genome was estimated to be between 106 (3) and 122 Mb (43). The assembly of TCC reported here consists in a “diploid” genome of 86.7Mb. However, the total assembly length is virtually the same if only contigs longer than 10 kb are considered (85.5 Mb). This value is below the expected genome (diploid) size, if one assumes that it should be similar to CL Brener, yet. it represents a very significant improvement compared to the best previous assembly obtained by Weatherly *et al.* (2009), whereby both CL Brener haplotypes combined (Esmeraldo like and Non-Esmeraldo) have a total added size of only 54 Mb (9). The smaller size of our assembly can be attributed to two factors. First, like CL Brener, TCC is a hybrid clone composed of two relatively divergent parental lineages similar to Esmeraldo and Non-Esmeraldo. This would imply that the distinction and “segregation” of parental haplotypes was partial (about 40% appears to have remained un-separated). It is likely that some genomic regions exhibit interallelic (inter-haplotype) divergence below the identity threshold that the assembler requires to discriminate between them. Secondly, it is possible that some clusters of repetitive sequences, especially the largest ones, were not completely uncollapsed, thus contributing to assembly shortening. It is difficult to estimate the impact of this second source of uncertainty on the assembly, since in the case of Dm28c, in spite of repetitions, the assembly was not smaller than the expected genome size. Importantly, however, for TCC we were able to distinguish and assemble separately (to a substantial degree) the parental haplotypes (Fig. 3). In fact, the dissection of the genomes into their constituent haploid genomic blocks is important to study some aspects of the evolutionary process that have been taking place in *T. cruzi* and could yield novel insight into the generation of hybrids and their subsequent evolution. An opportunity offered by our data, is the ability to detect events of homologous recombination between haplotypes. With this in mind, we mapped Illumina reads from Esmeraldo (DTU II) (download from SRA: WUGSC SRX271443) to the TCC genome. The two haplotypes exhibit a divergence of 4% on average, so reads of 75 nt in length are expected to have more than two mismatches with non-Esmeraldo haplotype. This analysis was performed with high stringency (allowing no mismatches) so that reads will basically map only to the Esmeraldo-like haplotype. The separation was evidenced by the read mapping distribution: almost all of them mapped only to one of the homologous contigs (Fig. 3(b)). However, in sites where recombination presumbaly took place, there is a switch in the mapping pattern and reads start to map on the homolog contig, as shown in Fig. 3(b). To confirm that the switch in the mapping preference was not an assembly artifact; namely a chimera generated by mixing both haplotypes, we mapped Pacbio reads that span the region of recombination to these sites (Fig. 3(c)). As shown in Fig. 3(c), mismatches between reads and the contigs are evenly and randomly distributed along the genome. In contrast, when mapped into chimerical assembled contigs (Fig S1), PacBio reads have low or high abundance of mismatches before and after the artificial “recombination” point. The mapping pattern of Illumina reads onto chimerical contigs is, nevertheless, indistinguishable from that observed in *bona fide* recombination zones.

**Fig. 3.**
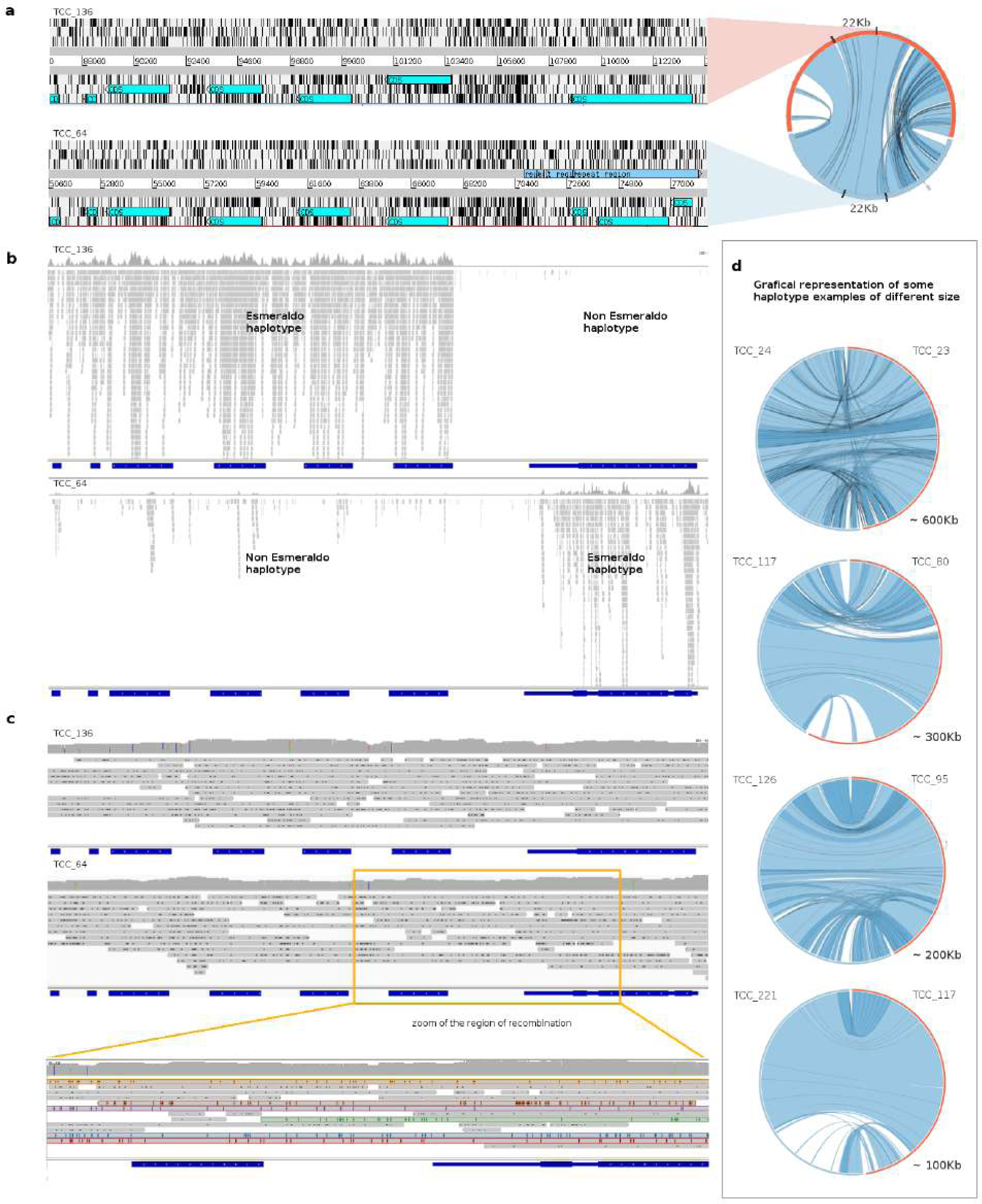
Haplotypes resolution and recombination. **A)** Circos graph representation of homologous contigs (right). On the left is shown the Artemis view of the indicated fragments (for contig TCC_133 from 88Kb to 112Kb (top), and for contig TCC_64 from 50Kb to 77Kb (botton)). The six frames are shown and the annotated genes are represented in turquoise. **B)** alignment visualization (IGV) of the Esmeraldo illumina reads (SRA833800) on the same homologous regions considered in **A** (TCC_133 on the top, TCC_64 on the botton). **C)** Alignment visualization (IGV) of PacBio TCC reads on the same region as in **B**. On the botton is represented the enlargement of the boxed region where Esmeraldo illumina reads go from mapping to TCC_133 to mapping to TCC_64. **D)** Circos graph representation of haplotype resolution contigs of different sizes.

Another noteworthy observation from Fig. 3b is an extreme scarcity of Illumina reads mapping on this putative recombination hotspot. To investigate this we tested several possible causes. First, we wondered whether this could be the result of our stringent mapping conditions. However, relaxing mapping restrictions to allow more mismatches only caused a marginal increase in mapping. The same is true for 454 reads; very few of them map here, even when a large percentage of mismatches is tolerated (from NCBI PRJNA50493). To us, this suggested that this region is either refractory to sequencing or an artifact of our assembly. To test the latter, we searched for the presence of the region in other assemblies and strains. It was found in Dm28c (our assembly), Sylvio X10/1 (from NCBI PRJNA40815, PacBio) and CL Brener (Sanger). This led us to conclude that this region is poorly sequenced using other NGS technologies. Further analysis of the region, revealed a poor GC content, averaging 30%, (with segments longer than 100 nt with GC content lower than 20%). This is in keeping with previous reports showing that genomic segments with biased low GC content are underrepresented in Illumina sequencing (44). Using GC content and Illumina sequencing depth as criteria for identifying similar regions in the genome of TCC, we found more than 500 regions simultaneously matching these two features: very low GC content and very few or none mapping reads (with segments where sequencing depth is as low as zero). A similar search in Dm28c (PacBio assembly), yielded 160 such regions. Comparing with other assemblies we found that most GC-low regions from TCC are also present in CL Brener assembly, but less than one half of them can be found in our assembly of Dm28c. The comparison of two Dm28c assemblies reveals that only 90 of the 160 found in the PacBio assembly, reported here, were present in the Dm28c assembly obtained using Roche 454 reads. Similar figures are obtained when the comparison is performed with Sylvio X10/1 (Illumina and Roche 454). Furthermore, most of these regions, or fragments of them, are located in contigs ends (for Illumina and Roche 454) indicating that assembly was halted in these positions due to insufficient coverage. Taken together, these results indicate that these GC-low regions are another important factor of assembly fragmentation in *T. cruzi* genomes for Illumina and 454 technologies, but they do not appear to affect Sanger or PacBio based assemblies.

### Genome annotation

Genome annotation is an error-prone task, which is particularly intricate in *T. cruzi,* due to its intrinsic genomic complexities including large multigene families, pseudogenization and the absence of fully annotated genomes from a phylogenetically closely related species. To address these problems, we followed an annotation strategy designed to handle the peculiarities of this species. Since the vast majority of trypanosome genes lack introns, we used a “prokaryotic-like” approach to identify them. Specifically, as gene models to be used downstream in the annotation pipeline we took a greedy criterion: all open reading frames (ORFs) longer than 450 nt and starting with Met. Shorter CDSs were dealt with separately (see below). Since many long ORFs are not protein coding, this initial group of potential protein coding genes very likely contains a large number of false positives. However only those ORFs encoding proteins will yield blastp hits with relatively distant species, hence, these can be readily identified and excluded by subsequent filtering with BlastP, using appropriate protein reference data sets. Therefore these data sets are crucial to get accurate annotations. Indeed a common drawback in genome annotation is inheriting (by homology transfer of information) spurious and erroneous annotations from others genomes used as references. To work around this potential source of inaccuracy the following dataset were built/used: i) A carefully curated databases of *Trypanosome cruzi* multigene families (TS, mucin, MASP, GP63, TASV, DGF-1, RHS). Each of these families was thoroughly scrutinized manually to determine the complete and probable functional copies, their sub-groups and pseudogenes. ii) Trypanosomatid proteome databases from which *T. cruzi* strains as well as other species not very evolutionary distant from *T. cruzi* (*Trypanoma rangeli, T. grayi*) were excluded. The rationale for this criterion is that whereas evolutionary conservation is an indication of functional relevanc*e (i.e.* if the amino acid sequence encoded by the ORF has been maintained over time it suggests a real protein coding gene), conservation among not distant taxa does not guarantee functional relevance; instead, it could merely reflect phylogenetic inertia (*i.e.* not enough time to diverge). iii) Curated protein database from non-trypanosomatids iv) Additionally, to cross-check our annotations, we used protein annotations from *T. cruzi,* exclusively from CL Brener, since this is the strain most accurately annotated. Whenever inconsistencies arose, problematic ORFs were further analyzed and manually curated. An overview of the workflow is presented in Fig. S2.

In order to include, and annotate protein coding genes shorter than 450 nt, the search was conducted backwards; namely a dataset of short proteins was build and used to search for them (using tblasn) in the assemblies. A battery of diverse tools was used to incorporate additional information in the annotation. Non-coding RNAs, transposable elements and repetitive elements, were annotated using dedicated software run on specific databases that were built for these purposes.

Although this strategy is intended to minimize misannotations, further manual curation were needed, specially with genes belonging to multigene families and their associated pseudogenes. Particularly, several of these genes were frequently excluded from previous assemblies, as they were impossible to accurately position (9). The genomes of the two strains presented here contain 27522 and 17371 genes (TCC and Dm28c respectively) and the number of hypothetical genes have been reduced from ˜50% to ˜39% by means of the removal of spurious annotation due to the inclusion of new genes functions that could be classified due to the improvements of the TriTrypDB, or because they were new coding sequences that were not assembled before.

To visualize and comfortably handle the information of *T. cruzi* genomes we generated a web platform (http://bioinformatica.fcien.edu.uy/cruzi/). A genome browser can be used to navigate the annotation of contigs longer than 50 Kb. It also offers built in tools to select group of genes and other genomic features that are distinctive of *T. cruzi* such as surface multigene families, repetitive elements and directional gene clusters. Among other functionalities of the interface, it is possible to visualize gene annotations and retrieve their nucleotide and amino acid sequences. It is also possible, to conduct different searches (by annotation or by keywords) and to visualize graphical representation of repetitions and haplotypes (haplotype information is available). The interface is intuitive and user friendly. See materials and methods for further details.

### The genome of *T. cruzi* is compartmentalized

Two groups of protein coding genes were very well described in *T. cruzi*: the multigene families with hundred of copies: DGF-1, GP63, MASP, mucins, RHS and TS, and those generically defined as "conserved", which can be further divided into two groups: genes encoding for proteins with a known function called “conserved genes”, and genes without an assigned function but present in more than one trypanosomatid species called “hypothetical conserved genes”. We confrimed that the genome of *T. cruzi* is compartmentalized in two clearly defined regions: a "core compartment" composed of conserved and hypothetical conserved genes, and a “non-syntenic disruptive compartment" composed of the multigene families TS, MASP and Mucins (Fig. 4(a)). On the other hand, GP63, DGF-1 and RHS multigene families have a dispersed distribution in the genome comprising both compartments, which may in turn be organized as unique or in tandem array distribution. Since the “core compartment” corresponds to the previously described syntenic blocks in *T. brucei* and *L. major* (4), and the “disruptive compartment” is mainly composed by species or genus specific genes, the latter can be considered as a recent region of the genome. It is noteworthy that the members of the disruptive compartment have been previously referred to as sub-telomeric; however, as it is evident from our results, these genes can be located in any position of the chromosome with the only condition that they cover wide ranges of distances, and the location can be anything from internal chromosomal regions to extremes of chromosomes, and even comprise whole chromosomes. Therefore the term sub-telomeric, probably inherited from the genome organization of VSG genes in *T. brucei*, is a misleading term that can lead to confusion. We suggest it should be replaced by the more encompassing concept of “compartments”.

**Fig. 4.**
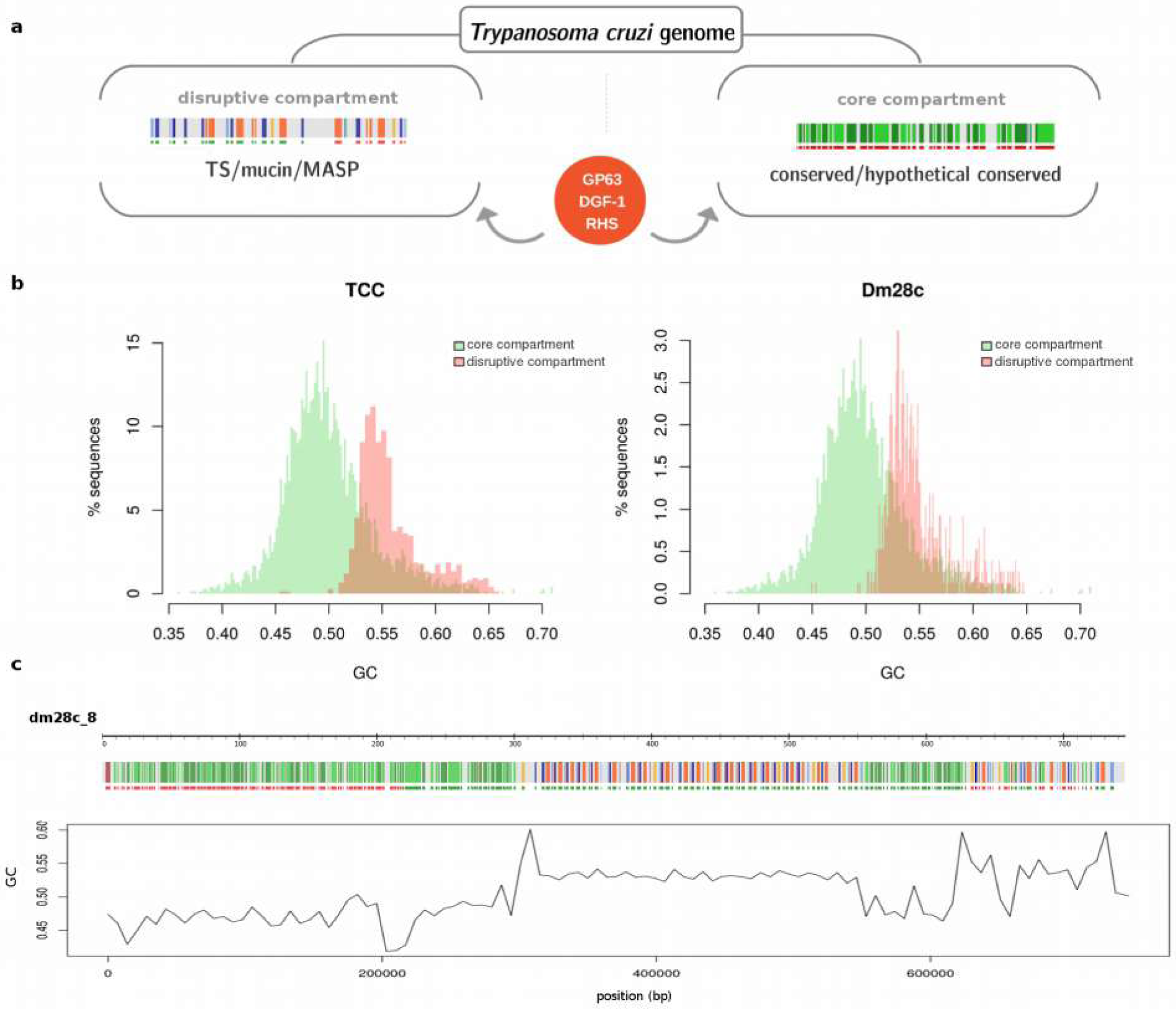
The genome compartmentalization of *T. cruzi*. A) Schematic representation of both type of compartment in *T. cruzi*. Genes are visualized as in the web interface by strips (DGF1 in red, GP63 orange, MASP blue, mucin light blue, TS light orange, conserved genes in green). The core compartment is composed by conserved genes. The disruptive compartment is composed by surface multigene families. GP63, DGF-1 and RHS although they are highly present in the disruptive compartment also are distributed-sometimes in tandem clusters- in the core compartment. B) GC distribution of the compartments. Only contigs entirely composed by one compartment (80% of higher proportion of conserved genes or surface multigene families) and longer than 10Kb were considered. C) Schematic representation of a contig of Dm28c (as in the web interface) and the GC distribution over a sliding windows of 7000bp.

Since compartmentalization is evident in both sequenced genomes, it probably constitutes a property of *T. cruzi.* Therefore, we aimed to identify general features distinguishing said compartments. For this, we performed compositional analysis of the genome, and found that the disruptive compartment consistently exhibits higher GC content than the core compartment (Fig. 4(b)). The bias in GC content becomes clear by simply plotting the GC distribution over the length of a contig as shown in Fig. 4(c). This genome organization resembles isochore-like structures. In fact, although originally only described in vertebrates (45,46), these organization elements were recently proposed to be a feature of all eukaryotes including unicellular ones (47,48). Our observations are consistent with this concept. This distribution deserves further studies to understand how compartmentalization emerged and evolved.

### Multigene families

One of the known characteristics of the *T. cruzi* genome is its widely expanded content mainly due to the large number of multigene families (49–51). In fact, when the first genome of *T. cruzi* was sequenced, authors warmed about the risk of underestimating the number of units of genes organized in tandem (3). Later, Arner and collaborators exposed examples of copy number underestimation and misassembly, and proposed that the number of protein coding genes and pseudogenes may be twice the previous estimates (52). Our genome assembly makes it possible to determine the real extent of the *T. cruzi* gene expansion. Defining a gene family as that which presents eight or more genes, we identified 90 and 74 families in TCC and Dm28c, respectively. A more detailed analysis of families by MCL clustering shows that they can be subdivided; TCC presents 190 paralogous gene clusters and Dm28c presents 139 ones (Table S2). In Table 2 we show only those gene families with more than 50 members in at least one sequenced genome. As previously described, the most enriched ones are TS, MASP, RHS, Mucins, DGF-1 and GP63. Remarkably, whereas RHS, DGF-1, GP63 and TS fall into 6 or less clusters in both strains, mucins, and particularly MASPs proteins, harbor higher number of subfamilies (clusters) (Table 2 and Table S2). On the other hand, misannotation of mucins and TS led us to a more in detail analysis (see below). Interestingly, a more accurate estimation of copy number, and the analysis of the core compartment, allowed us to identify more than 20 families with at least 50 genes, including several genes defined as "housekeeping" in eukaryotes, such as Hsp70, Histones H2B and H3 (Table 2, Table S2). Given that gene expression is regulated mainly at the post-transcriptional level in trypanosomatids, the increase in the number of genes has been proposed as a mechanism for increasing gene expression level. However by gene ontology analysis we could not find any particularly enriched metabolic pathway which could further strengthen this hypothesis from a functional perspectives.

**Table 2.** Gene families groups in *T. cruzi.* The total number of genes and clusters are listed

| Gene Product | TCC* |  | DM28c* |  |
| --- | --- | --- | --- | --- |
|  | Members | MCL clusters | Members | MCL clusters |
| *trans-sialidase (TS)* | 1729 | 2 | 1489 | 3 |
| *MASP* | 1204 | 44 | 935 | 38 |
| *RHS* | 1264 | 4 | 774 | 2 |
| *Mucins* | 787 | 21 | 427 | 10 |
| *GP63* | 712 | 6 | 390 | 6 |
| *DGF-1* | 491 | 1 | 215 | 1 |
| *UDP-Gal or UDP-GlcNAc-dependent glycosyltransfer* | 128 | 1 | 110 | 1 |
| *Protein kinase* | 152 | 6 | 118 | 4 |
| *Amino acid permease/ transporter* | 128 | 1 | 93 | 1 |
| *Elongation factor (1-alpha, 1-gamma and 2)* | 109 | 3 | 63 | 3 |
| *Protein Associated with Differentiation* | 98 | 1 | 69 | 1 |
| *Glutamamyl carboxypeptidase* | 96 | 1 | 80 | 1 |
| *Syntaxin binding protein* | 91 | 2 | 40 | 2 |
| *TASV* | 87 | 5 | 53 | 3 |
| *Heat shock protein 70* | 87 | 2 | 40 | 1 |
| *Kinesin* | 85 | 3 | 55 | 3 |
| *Glycine dehydrogenase* | 73 | 2 | 51 | 1 |
| *Beta galactofuranosyl glycosyltransferase* | 73 | 1 | 37 | 1 |
| *Receptor-type adenylate cyclase* | 63 | 1 | 27 | 1 |
| *Histone H2B* | 56 | 1 | 5 | 1 |
| *Histone H3* | 54 | 1 | 27 | 1 |
| *Oligosaccharyl transferase* | 52 | 1 | 3 | 1 |
| *Tryptophanyl-tRNA synthetase* | 52 | 1 | 36 | 1 |
| *ATP-dependent DEAD/H RNA helicase* | 59 | 3 | 39 | 3 |
| *Casein kinase* | 50 | 1 | 42 | 1 |
| *Cysteine proteinase* | 46 | 1 | 56 | 1 |
| *Flagellar calcium-binding 24 kDa protein* | 41 | 1 | 52 | 1 |

Clusterization analysis also revealed specific clusters of hypothetical protein coding genes. We identified 47 (TCC) and 27 (Dm28c) novel multigene families of at least 8 paralogues, with 10 of this clusters containing 20 or more paralogues on each genome (Table S3). Among them we found shared and strain specific families containing conserved domains (Table S3). Remarkably, the largest family is composed by 158 members in TCC (cluster 35, Table S3) meaning that, after excluding the multigene superfamilies, it represents one of the most enriched genes in the whole of the TCC genome (Table 2), suggesting a relevant role in the parasite that deserves further study.

Interestingly, clusterizations include all the genes from a family, both in tandem and dispersed. In fact, although tandems of genes have been already described in *T. cruzi*, the resolution of previously collapsed repetitive regions allowed us to visualize and measure the extent of the tandem arrays of genes; we identified 2363 (TCC) and 1619 (Dm28c) genes organized in tandems of at least two genes (excluding hypothetical genes).This organization, which was previously difficult to identify, is now evident by simply looking at the chromosomes (Fig. 5(a) and Web interface). Needless to say that tandem genes were grossly underestimated in previous assemblies. Fig. 5(b) shows the increment on the number of tandem arrayed groups of genes, in comparison with the original CL Brener genome assembly; there are five times more gene tandems of 4 genes in TCC than those identified previously.

**Fig. 5.**
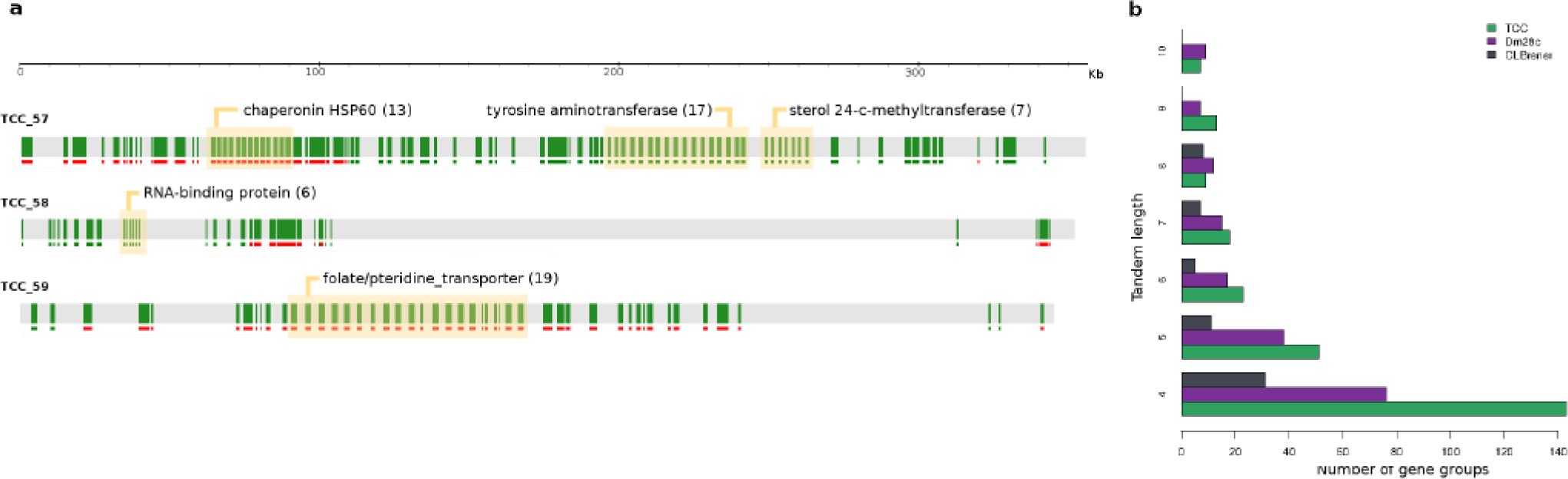
Tandem gene organization. A) Representation of three contigs of TCC as in the web interface where only conserved genes are shown (green strips). Groups of tandemly arrayed genes are highlighted. B) Graph representation of the number of groups of tandemly arreayed genes (represented tandem length from 4 to 10 genes) in the different genome assemblies. TCC in green, Dm28c in violet, CL Brener in gray.

### Single copy genes

Although multigene families encoding for surface proteins are the main cause of genome expansion, this feature is a more general phenomena of *T. cruzi*, affecting the majority of its genome content. In fact, most genes have at least two copies per haploid genome. Accuracy in gene copy number determination is highly relevant for functional studies, since knock out experiments and the generation of transgenic strains constitute an essential source of information to determine gene essentially, paramount to defining new drug targets for treatment of Chagas disease. Keeping this in mind, we wondered what the actual core of single copy genes was. We found approximately 5000 single copy genes in Dm28c, of which only 1377 have an assigned function. We then focused our analysis in those genes whereby knock-out experiments were reported (Table S4). First we noticed that a number of genes previously considered to be single copy form a cluster of a few number of copies, which led to miscalculations when semi-quantitative methods were used. Such is the case for lipoamide-dehydrogenase, whereby the copy number was determined by Southern blot (53). This gene exhibits seven copies in TCC and six in Dm28c. *msh2* (54) has four and two copies in TCC and Dm28c, respectively, and *hgprt (55)* presents four copies on each genome. *T. cruzi* has been traditionally considered an organism refractory to genetic manipulation. The difficulties to perform gene knock outs (*e.g.* to insert genes of resistance to a drug with 5’ and 3’ flanking regions of the target gene) has been ascribed to particular features of its recombination machinery. However, further analysis of the few successful KO experiments reported (56) revealed that all of them (*gp72, lyt-1, kap3)* exhibit unique copies in both strains (57–59). We propose that copy number underestimation could account for the failure of classical knock-out experiments in this parasite. Conversely, single copy genes that could not be completely knocked out such as calreticulin, *dhfr-ts* and *cub*, but instead monoallelic deletions were obtained should be considered essential (60–62). Thus, long read sequencing allowed correct estimation of copy number, leading us to reconsider the assumption that classical gene KO methods do not work in *T. cruzi*. On the other hand, our data can also aid in predicting the feasibility of mutagenesis experiments done using CRISPR technology whereby the higher the copy number, the higher the difficulty to “hit” all copies simultaneously.

### Mucins and trans-sialidases

The biological relevance of mucins and trans-sialidases in the infection, in addition to the high number of incomplete genes and imprecise pseudogene number determination, motivated us to manually curate these genes.

Mucin-like proteins of *T. cruzi* were classified in three main groups, TcMUCI, TcMUCII, and large and small TcSMUG (TcSMUGL and TcSMUGS respectively). TcMUCI and TcMUCII are expressed in the mammalian stages of the parasite, having a highly variable region and molecular weights ranging from 80 to 200 kDa (51,63), whereas SMUG genes, smaller than TcMUC and with little variability, are mainly expressed in the insect derived stages (50,64,65). Initially classified as TcMUCI and TcMUCII, the presence of a mosaic of sequences intermediate between both groups led to the proposal of a common ancestor and further diversification (51). In this work we considered as complete mucin genes those whose deduced amino acid sequences had an N-terminal signal peptide (SP), a C-terminal GPI anchor sequence, and T-rich sequences such as T8KP2, T6-8KAP or T6-8QAP. With these criteria and including RNAseq data (31,32), we performed manual searches, including correction of initial methionine in some cases. In this manner, we identified a total of 970 and 571 mucin genes in TCC and Dm28c, respectively, of which 247 (TCC) and 113 (Dm28c) correspond to pseudogenes (Table 3). Comparison between both strains led to practically the same profile of mucin groups; around 60% were classified as TcMUCII and only 6% as TcMUCI. However, we found differences in the distribution of the TcSMUG family. In TCC ˜6% are TcSMUGS and ˜3% TcSMUGL while the opposite occurs in Dm28c where ˜4% were TcSMUGS and ˜10% TcSMUGL. We found 56 genes coding TcSMUGL in Dm28c and only 25 in TCC. These findings raise the interesting possibility that these differences may be associated to phenotypic differences in virulence, or related to different DTUs, and remains to be investigated. Our analysis also revealed the presence of a large number of mucins fragments and/or pseudogenes, in a similar percentage for both strains: 25 and 20 % in TCC and Dm28c, respectively.

**Table 3.** Manually annotated surface multigene families

|  | TCC |  |  | Dm28c |  |  |
| --- | --- | --- | --- | --- | --- | --- |
|  | total | genes | pseudogenes | total | genes | pseudogenes |
| TS | 1734 | 1013 | 721 | 1491 | 924 | 567 |
| MASP | 1332 | 888 | 444 | 1045 | 666 | 379 |
| GP63 | 756 | 443 | 313 | 416 | 267 | 149 |
| DGF-1 | 491 | 207 | 284 | 215 | 83 | 132 |
| **Mucin** | 970 | 723 | 247 | 571 | 458 | 113 |

| Mucin |  | TcMUCII | TcMUCI | TcSMUGS | TcSMUGL | pseudogenes |
| --- | --- | --- | --- | --- | --- | --- |
| TCC | 970 (100%) | 585 (60.3%) | 57 (5.9%) | 56 (5.8%) | 25 (2.6%) | 247 (25.5%) |
| Dm28c | 571 (100%) | 351 (61.5%) | 29 (5.1%) | 22 (3.9%) | 56 (9.8%) | 113 (19.8%) |

Trans-sialidase/trans-sialidase like proteins (TS) constitute a large and polymorphic superfamily of around 1400 members. This heterogeneous family is currently classified in eight different groups, and although most of them do not exhibit trans-sialidase activity they play relevant roles in infectivity and virulence (66). TS present an N-terminal SP, and a C-terminal GPI anchor signal. Functional members of TS family present the characteristic domain VTVxNVLLYNR. Additionally, some groups have N-terminal ASP box sequences. Groups I and IV also present SAPA repeats, an extremely antigenic region whose role is to increase the half-life of TS in blood (67). The initial annotated TS were manually filtered using the following criteria: presence of VTV domain, presence of ASP box domain, GPI anchor signal probability and presence of a SP sequence. Using these criteria we found 1734 TS genes in TCC and 1491 genes in Dm28c. Our analysis also revealed the presence of a large number of pseudogenes or fragments; 721 and 567 for TCC and Dm28c, respectively (Table 3).These results highlight the importance of manual curation for the most expanded and complex multigene families.

### Transposable elements

Transposable elements are dynamic drivers of evolutionary processes that contribute to genomic plasticity. Usually present as repetitive sequences in the *T. cruzi* genome, they are abundant and commonly misannotated. *T. cruzi* presents three families of autonomous genomic elements, as well as their non-autonomous pairs. Here, we were able to identify the entire and their flanking-sequences of all TEs families present in both genomes (Table 4). Namely, VIPER, a tyrosine recombinase (YR) element which belongs to the DIRS order; L1Tc, a non-LTR element of the INGI clade; and CZAR, also a non-LTR element from the CRE clade which is site-specific inserting only on the SL gene (3,68– 70). On the other hand, non-autonomous elements have also been identified. Namely, SIDER which has sequence similarity to VIPER´s 5’ and 3’ ends, NARTc, the non-autonomous pair of L1Tc elements, and TcTREZO (71). Putative active and defective copies could be determined for all families. None of the VIPER, CZAR and TcTREZO elements had complete domains, indicating that all copies are defective, whereas L1Tc was the only one to show putative active copies in both genomes.

**Table 4.**
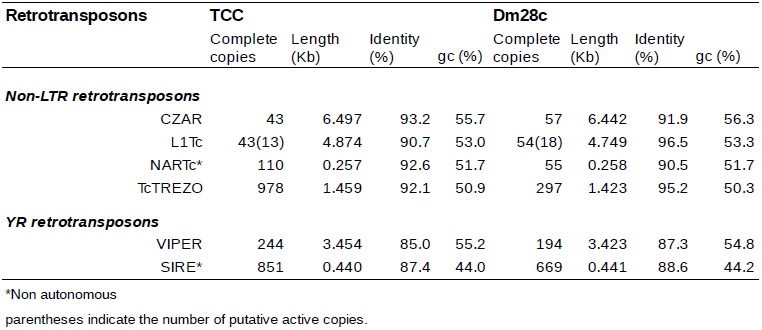
Complete retrotransposon copy numbers in *T. cruzi.*

To shed some light on the evolutionary dynamics of L1Tc transposon, we first built a maximum likelihood phylogenetic tree using only complete, and hence putatively functional sequences of this element. The resulting tree (Fig. 6), exhibits three main clades, one containing L1Tc copies from Dm28c, and two clades containing TCC copies. To further explore this configuration, we analyzed (full length) L1Tc elements from CL Brener, discriminating Esmeraldo and Non-Esmeraldo haplotypes. It becomes immediately apparent that while one TCC cluster is associated with Esmeraldo haplotype, the second TCC cluster is related to non-Esmeraldo haplotype (Fig. 6). This evidences that the functional copies of L1Tc had significant activity and underwent substantial evolutionary divergence after the two ancestral *T. cruzi* lineages conforming TCC (and CL Brener) split apart. The fact that branches in these two clusters are considerably long, denotes a low transposition/divergence ratio. It is surprising, however, that there are hardly any intermingled copies, meaning that any inter-copy variability predating the separation of these two ancestral lineages (i.e. already present in the ancestral genome) has been erased. This is consistent with the fact that Dm28c copies are also isolated forming a single cluster. Nevertheless, in this case most copies are very similar to each other (short branches) suggesting that in this strain, L1Tc exhibits a much higher transposition/divergence ratio. One can attribute this to the fact that the new copies did not have enough time to diverge after their emergence owing to high transposition rate. Alternatively, this might be due to a much slower nucleotide evolutionary (divergence) rate. Incorporating in the analysis the set of L1Tc sequences from the strain Sylvio X10/1 (from NCBI PRJNA40815) gives some clues on the cause of the different dynamics. In effect, these sequences cluster together with those of Dm28c, yet a sharp division between both strains is observed (Fig. 6). Since the two strains are very close relatives (with virtually identical nucleotide sequencies genomes wide), short branches in this part of the tree are cannot be attributed to slow divergence rate, favoring the hypothesis of very high transposition rate in this DTU. In any case, it denotes a markedly different dynamic from that observed in the hybrid DTU VI. Whether the different dynamics are related to the hybrid/non-hybrid nature of the strains being considered is a subject that deserves further assessment.

**Fig. 6.**
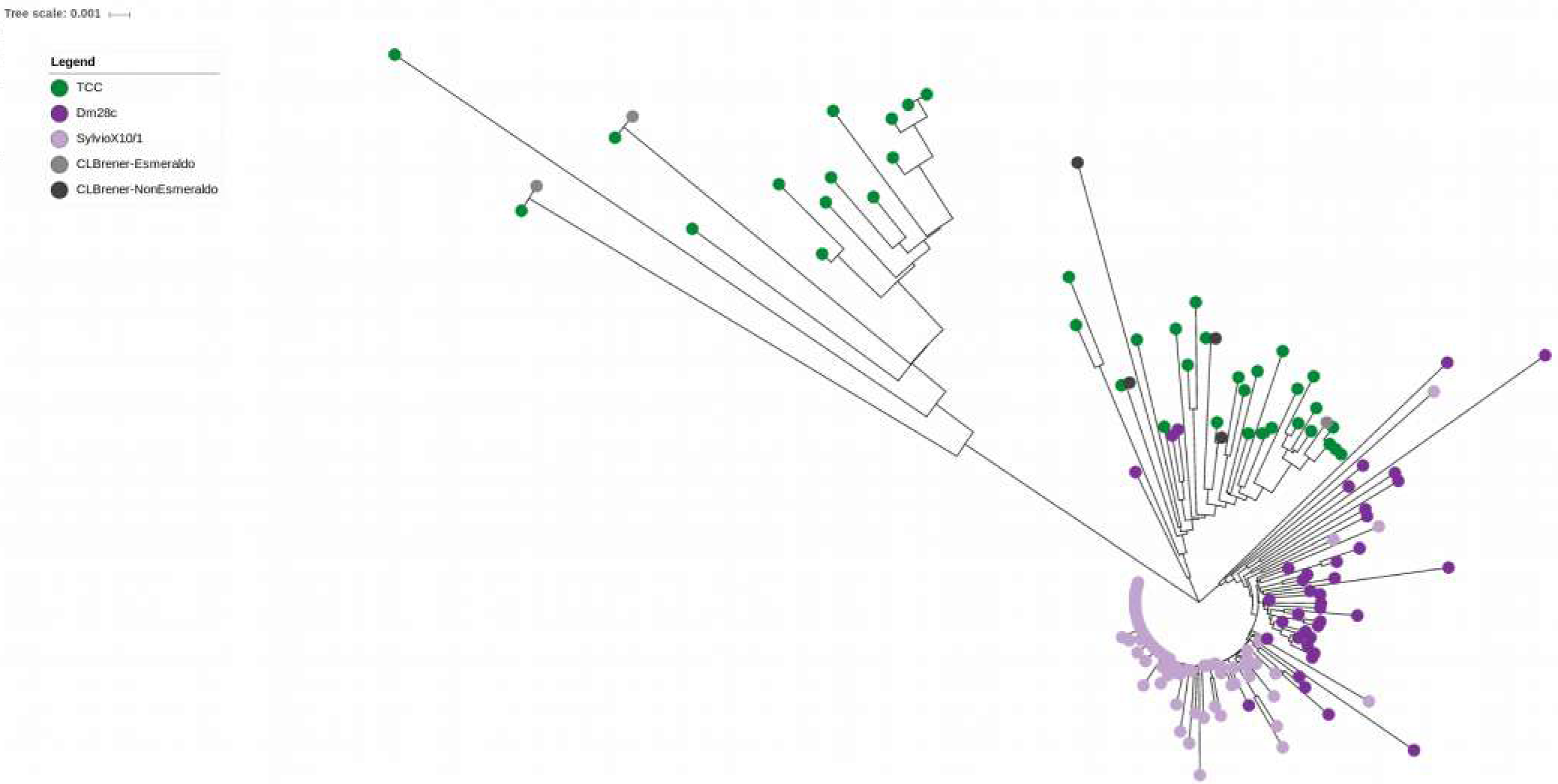
L1Tc phylogeny. Maximum likelihood phylogeny of complete sequences of L1Tc. Elements form TCC in green, Dm28c in violet, Sylvio (from PRJNA40815), CL Brener Esmeraldo like in light grey, CL Brener NonEesmeraldo like in grey.

### Tandem repeats

Tandem repeats (TR) are considered "neglected" sequences in genome analyses, since short reads cause often unsolvable problems for *de novo* assemblies in TR regions. As a consequence of long-reads we could resolve TR enriched regions previously fragmented. As expected, thousands of copies of the already well characterized 195nt satellite (72,73) were found: 41061 (8,3Mb) and 12244 (2,5Mb) copies for TCC and Dm28c, respectively (Table 5). These findings are in keeping with previous estimates made by cell and molecular biology methods (71,74). The difference in copy number between the genomes analyzed are not a consequence of the different genome sizes only. In fact, it has been observed that satellite amounts are variable among strains, reaching differences of up to six fold, but organized similarly throughout the genome (74). Our analysis shows that the satellite is two times more abundant in TCC than in Dm28c, accounting for 9.5% and 4.7 % of the total size of the genomes, respectively (Table 5). This coincides with the differences estimated between TcVI and TcI groups by Souza et al, being the repeats in the former 2-4 times more abundant than in the latter. Additionally, we recovered satellites reaching 45Kb and 42Kb in Tcc and Dm28c, but the majority were of smaller size, with an average of more than 10Kb, as previously reported (74). Interestingly, we could assembly the sequence around some satellite, and confirm the vicinity of the satellite clusters (Fig. S3).

**Table 5.** Annotated tandem repeats in TCC and Dm28c

|  | satellite | # tandems | Total Length (bp) | % Genome | Mean Id (%) |
| --- | --- | --- | --- | --- | --- |
| TCC | 195bp | 41062 | 8,303,881 | 9.5 | 95.4 |
| Dm28c | 195bp | 12244 | 2,483,120 | 4.7 | 95.1 |

|  | Tandem length | # groups | Total Length (bp) | Mean Id (%) | Belongs or include genes (%) |
| --- | --- | --- | --- | --- | --- |
| **TCC** | 3 to 10 | 74 | 154,516 | 85.5 | 39.2 |
|  | 11 to 100 | 135 | 833,791 | 84.2 | 74.1 |
|  | 101 to 500 | 26 | 343,243 | 94.7 | 80.8 |
|  | 501 to 2000 | 26 | 1,528,929 | 97.6 | 92.3 |
| **Dm28c** | 3 to 10 | 59 | 117,028 | 85.0 | 37.3 |
|  | 11 to 100 | 104 | 185,816 | 84.1 | 62.5 |
|  | 101 to 500 | 18 | 301,938 | 96.0 | 66.7 |
|  | 501 to 2000 | 24 | 1,091,688 | 97.9 | 91.7 |

We then aimed to characterize the remaining TR sequences with a minimal length of 3nt and containing at least 10 monomers. We identified 261 and 205 groups of different sizes being ˜1960bp the longest period in both genomes (Table 5).

The identified TR (related or not to coding sequences) identified here are shared between the strains analyzed. Around 85% of TR are present in both genomes, although only 107 groups of TR are shared. This means that the remaining, even when they are present in the other genome, do not meet the selection criteria; they do not have at least 10 copies or enough identity between copies (*i.e.* they are more degenerate in one genome than in the other one).

It is noteworthy that several TR are composed by protein coding genes organized in tandems where the intergenic regions are highly conserved. Such is the case of Flagellar Attachment Zone protein 1, R27-2 protein, Kinesin-like protein and histone H4 among others (Table S5). This can represent recent events of consecutive gene duplication that, in contrast with the tandem arrays of protein coding genes described above, still retain high levels of identity among their intergenic regions. In fact, around 90% of the TR of period from 500 to 2000 bp corresponds to these arrays, and the tandem intra-identity is above 97.5% (Table S6). Conversely, we detected TR of shorter periods present within coding sequences (Table S6). These internal tandem repeats are multiples of three, which indicates that they will be translated as a tandem of amino acids. This feature was first identified on the shed antigen SAPA (75), and subsequently, by immunoscreening of expression libraries (76,77). Additionally, TRs have been observed in TS, mucins and MASP, and in several other hypothetical proteins exhibiting signal sequences or trans-membrane domains, some of which were tested as antigenic (78). Antigenicity of these tandems, and their wide distribution along CDS suggest that they constitute a general mechanism for including unspecific polyclonal immune responses. These could act as "smoke screens", which could allow the parasite to evade the host immune response.

Overall our results highlight the power of improvement in whole genome studies offered by PacBio technology. Long-reads allowed us to accurately assemble, annotate and analyze the complex *Trypanosoma cruzi* genome, opening new perspectives in biological studies oriented towards the eradication of Chagas disease.

## Materials and methods

### Parasites and DNA isolation

*T. cruzi* strains Dm28c (12) and TCC (13), were used throughout this work. Epimastigotes were grown in liver infusion tryptose (LIT) medium supplemented with 10% Fetal Calf Serum at 28° C;total DNA was extracted using Quick DNA ^™^ Universal kit (Zymo Research, USA).

### Sequencing

PacBio library preparation and sequencing were done in the University of Washington PacBio Sequencing Services. Briefly, DNA was mechanically fragmented using a Covaris g-TUBE device, and concentrated with AMPure PB magnetic beads. The final long insert PacBio libraries were size selected for fragments larger than 10 kb using the BluePippin device.

A total of 8 SMRT cells were used, 5 to TCC and 3 to Dm28c yielding 751460 and 601168 raw reads respectively. Subreads were obtained using the SMRT Analysis RS.Subreads.1 pipeline (minimum polymerase read quality = 0.85); minimum polymerase read length and minimum subread length = 500 bp).Illumina MiSeq platform was used with paired-end library (2 × 150 cycles). Briefly, Nextera XT (Illumina, USA) library preparation kit was used from 1 ng of total DNA according to manufacturer instructions. Index primers were added to each library to allow sequence multiplexing. After 12 PCR cycles, the final library was purified with AMPure XP (Benchman, USA) and quantified with the Qubit dsDNA HS assay kit (Invitrogen, USA). Quality and length of the library were assessed with the Agilent high-sensitivity DNA kit (Agilent, USA) using the 2100 Bioanalyzer (Agilent, USA).

### Genomic assembly

The assembly was performed using the SMRT Analysis tools implemented in the HGAP pipeline V3 (14). It was run with the default parameters modifying only the expected genome size which was set to 110 Mb generating 1978 and 1143 contigs with an N50 of 73 Kb and 129 Kb for TCC and Dm28c respectively. Subsequently a scaffolding pipeline IPA was applied (15), specifically those scripts aimed to merge overlapping contigs using the assembled output and Illumina paired-end reads. This pipeline starts by self-mapping the contigs using megablast, with the parameter of word-size set at 40 and an e-value of 1e-80. Self-contained contigs and those with an identity below 99% and shorter than 500 bp were discarded from further analysis. Then Illumina reads were mapped over the remaining sequences using the alignment software SMALT. Finally, if there are enough pairs of Illumina reads where each member of the pair maps to two different contigs, it can be argued that the pair comes from the same contiguous sequence. This information was used to merge both contigs by their ends.

The final assemblies were of 86.7 Mb and 53.2 Mb with a N50 of 265 Kb and 318 Kb for TCC and Dm28c respectively. The assemblies were deposited at NCBI database BioProjects: PRJNA432753 and PRJNA433042.

### Genomic annotation

For the annotation of the coding sequences we extracted from each assembled genome the open reading frames of at least 150 amino acids of length, between a start and a stop codon, using the getorf tool from the EMBOSS suite (16). These sequences were mapped using blastp (17) against a database of kinetoplastid proteins from TritrypDB (http://tritrypdb.org) using as cutoff an e-value of 1e-10. The database of kinetoplastid proteins consists in a curated collection of CDS from various *Trypanosoma sp (excluding T. grayi* and *T. rangeli), Leishmania sp., Leptomonas sp., Crithidia sp.,* and *Endotrypanum sp..*

For practical purposes the annotation was divided into several steps. First the multigene families (MASP, mucin, RHS, GP63, trans-sialidase, DGF-1 and TASV) were annotated using a stricter cutoff, an e-value of 1e-30. Then we annotated the conserved genes among kinetoplastids. In this step we chose the best hit in *T. cruzi* CL-Brener Esmeraldo and NonEsmeraldo strains, and the best hit outside *T. cruzi*. If there were hits only outside *T. cruzi* these ORFs were also annotated with the best two hits. The selection of the best HSPs was performed using a personalized script that scanned for the lowest possible e-value with a meaningful description. If the only description available was ’hypothetical protein’ or a similar non-descriptive tag, then the lowest e-value was selected. For annotating short proteins, those smaller than 150 amino acids; we extracted the open reading frames between 50 and 150 amino acids generating a massive amount of sequences that were subsequently mapped only against a *T. cruzi* database by blastp searches (e-value 1e-03, identity > 80% and query coverage > 80%).

A curated non-coding RNAs and interspersed repeats database (from TritrypDB and NCBI) was used for the annotation of these elements. The sequences in these databases were mapped using blastn against the assembled contigs, then filtering the results and keeping only those hits which mapped more than an 80% of the length of the queried sequence with at least an 80% of identity. An overview of the workflow is presented in Fig. S2.

Customized scripts were written in Python (18), perl (19) and R (20). Alignments of short reads (illumina and 454 reads) were performed with Bowtie (21) and Rsubreads (from within R,(22)). Samtools utilities (23) were used to manipulate alignments and perform coverage analyses

### Within and between contig similarities

#### Dotplot view

A self-comparative mapping of each contig showing the results in a dotplot was performed with blastn. We used the YASS web server (24) with the default options to create these plots.

#### Circos view

To assess the similarities between contigs of each assembly, we run a blastn of all-against-all the contigs longer than 50 Kb. For practical purposes we generated two tables, one with HSPs longer than 5 Kb and other with HSPs between 3 Kb and 5 Kb. To graphically represent this data and avoid an entangled plot with an over-representation of hits from multigene families, the hits from these sequences were removed from the final results. The mappings between contigs are shown in a circular layout using the Circos software (25). In these plots (Fig. 3(d) and web interface) the target sequence is shown at 12 o’clock and extends clockwise proportional to its length. The ten contigs with the largest overall mappings were selected, and are represented according to their length in the plot. The genomic similarities of the target sequence and the contigs are represented as wide lines that show the length of the mappings, their coordinates, and share the same color with the contig of origin.

### The web interface

An online genome browser was developed (bioinformatica.fcien.edu.uy/cruzi) where the whole annotation is displayed for contigs longer than 50 Kb. Each contig is shown as a long rectangular block where genes and sequences of interest are represented as rectangles of different colors according to their classification. The width and position of each rectangle is proportional to their length and coordinates in the contig. Each represented contig has at the bottom a smaller rectangle that indicates the direction 5´-3´: sense (red) or antisense (green). In the visualization the annotation is divided in different categories. The CDSs are divided according to the main multigene family they belong, conserved genes are divided between those with a meaningful description and those annotated as hypothetical proteins. The repeated elements are also classified as tandem, interspersed repeats or satellite. Each category is shown with a different color and can be displayed or hidden as a whole from the check boxes in the upper part of the page. When the cursor is over an annotated sequence it displays a tooltip with a description of the best blast hits and hyperlinks to the nucleic and amino acidic sequence as long as possible. Under each contig there is a white rectangular block with more information. There are hyperlinks to the contig sequence in fasta format. There are also three plots, two that show how it mapped against other contigs in a circular plot, the leftmost shows HSPs longer than 5 Kb, and the one in the right the HSPs between 3 and 5 Kb. The right-most plot is a dotplot that represents how the contig maps against itself (see within and between contig similarities for more details). On the bottom-right of the page there are five buttons, two ones with the magnifying glass allows to zoom in and out in the sequences, the one with the stripes allows to change views between the standard view and a six frames view, and the text box and binoculars allows to search for sequence descriptions within the page. It is also possible to extract a customized nucleotide sequence from the contig by clicking at the start and end of the desired area. After doing this an emergent window appears at the bottom of the page allowing to correct the coordinates and retrieve the genomic sequence.

### Chromosomal and contig visualization

Besides the developed web interface we used ACT (26) and ARTEMIS (27) to visualize and compare contigs from our assembly and previous assemblies. Both software use the output of the blastn search and the contigs in fasta format. Blastn search for these comparisons were performed with stringent e value < 1e-200. IGV (28) software was used to visualize the alignments of reads over the assemblies (including illumina, 454 or PacBio reads of different sources)

### Manual curation of surface multigene families annotation

The annotation of surface multigene families was based on the coding sequences identified by the annotation pipeline. Further, we manually inspected the sequences analyzing specific characteristics relatives to their function.

For mucins we use the length, the presence of GPI, SP, the number of repetitions of threonine tandems, the presence of RAPS and TCRLL domains and the S,T and P amino acid composition. GPI anchor sites were predicted by predGPI (29) and signal peptide by SignalP v4.1 (30) taking in consideration that some genes carried a missannotated initial methionine. The identification of conserved N and C terminal domains as well as characteristic sequences and tandem repeats present in mucins were performed with custom scripts. Genes without a clear signal peptide and GPI anchor signal, but containing other mucin properties were designated as pseudogenes (most of them correspond to genes that lost the N or C terminal extremes). Finally, analysis of RNA seq data (31,32) and correction of initial methionine were performed where applicable.

For trans-sialidase sequences we considered the length, the presence of VTV and ASP box domain, the GPI anchor signal probability and the presence of a signal peptide sequence (same software than above). SXDXGXXTW, VTVxNVxLYNR and LYN domains were searched using PatMatch (33) and the output parsed with customized scripts.

For MASP genes, we consider the presence of highly conserved domains. N-terminal domain MAMMMTGRVLLVCALCVLWCG and C-terminaldomain GDSDGSTAVSHTTSPLLLLLVVACAAAAAVVAA were searched by HHsearch (34). Genes having only one domain were annotated as pseudogenes.

For DGF-1, genes smaller than 1000 bp were discarded, and smaller than 2000 bp were considered as pseudogenes. Fragmented DGF-1 were annotated as pseudogenes when partial ORFs were found with less than 2000 bp of distance. The start of the first fragment and the end of the last are the coordinates for the annotation.

### Gene cluster analyses

Protein-coding genes were clustered into gene families using MCL (35) with blastp-log E-values (e – 20). A fairly stringent inflation value which determines the granularity (or size of the output clusters) of 4 was used. A customized script was used to parse the clusterization output and generate the final results. Clusterization was performed to determine gene families, novel gene families and single copy genes.

### Characterization of transposable elements

Canonical complete elements for each TE were used to perform queries against the genomes using blastn (evalue 1e-10, identity > 80%, query coverage > 90%). Non autonomous elements (SIRE and NARTc) were filtered when mapped into the parental elements (VIPER and L1Tc). L1Tc was also identified in CLBrener assembly (from TritrypDB v28 Tcruzi CLBrener, Tcruzi CLBrener Esmeraldo-like and Tcruzi CLBrener Non-Esmeraldo-like) and Sylvio X10/1 assembly (from NCBI BioProject PRJNA40815) with same parameters. A total of 7 L1Tc element were identified in CLBrener and 156 in Sylvio X10/1. Multiple alignments were performed with MAFFT (36) using e-ins-i Iterative refinement method. Phylogenetic trees were computed by PhyML 3.1 (37) through see view plataform (38). A maximum likelihood tree based on GTR+i model was performed with 1000 bootstrap replications. Final representation was performed with iTOL (39).

### Characterization of tandem repeats

To identify and annotate tandem repeats we used the Tandem Repeats Finder software (40) with the parameters for minimum alignment score of 20, maximum period size of 2000, and the alignment score of 2 for a matching base and -7 for mismatches and indels. All entries with at least ten repeated periods of at least three nucleotides of length were considered for analysis. The results that share an overlap over 80 % in both sequences were merged and reported as one. The grouping of these sequences was done in several steps. First, we used personalized scripts to generate all the variants of a period and scan the results to group the identical ones. This strategy is better suited for grouping short periods without internal variability.

The second step of clusterization consisted in creating a multifasta file with a sequence from each period and grouping them using blastclust with the parameters -S80 -L0.8 -bT that restricts the results to those that share at least an 80% of the sequence length with and 80% of identity for both sequences. This grouping step is mainly aimed for clustering longer sequences with few indels or mismatches.

For further grouping, a third step was used, where each sequence was extended by repeating the period up to at least 300 bp. Then a self-comparative blast was performed and results with an identity lower than 80 % were discarded. Using a personalized script, for each sequence all the HSP were summed and if this sum was at least an 80% of the target sequence, they were grouped. This step is particularly useful for grouping sequences where the shortest reported period is in fact a multiple of a shorter one of another cluster. The final output is a table with the coordinates of each tandem repeat and the group they belong. An overview of the workflow is presented in Fig. S4

## Acknowledgements

We thank María Eugenia Francia and Lucía Spangenberg (Institut Pasteur de Montevideo) for critical reading of the manuscript, and Gonzalo Greif, Natalia Rego and Hugo Naya (Institut Pasteur de Montevideo) for technical assistance in bioinformatics and moolecular biology.

---

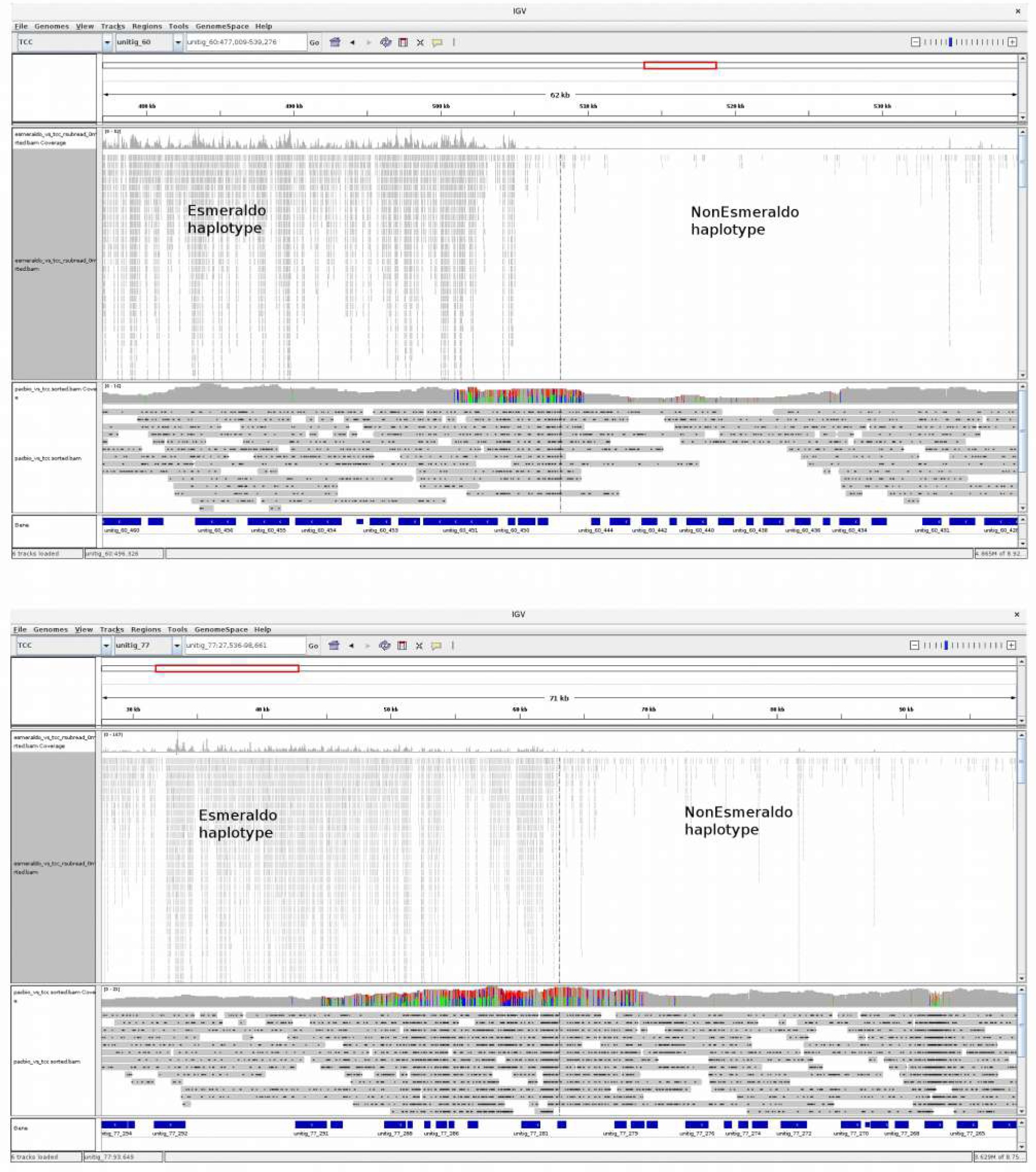

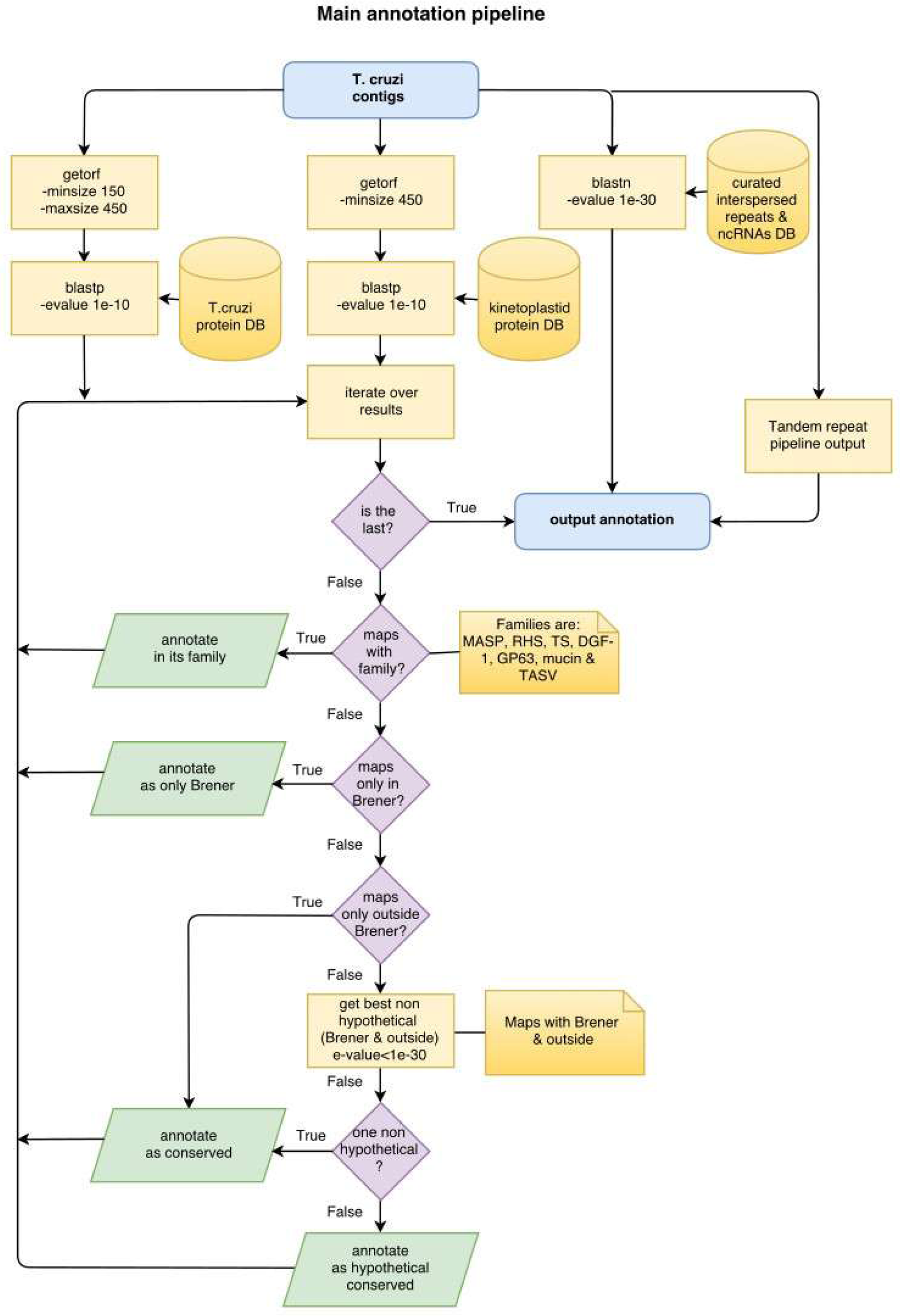

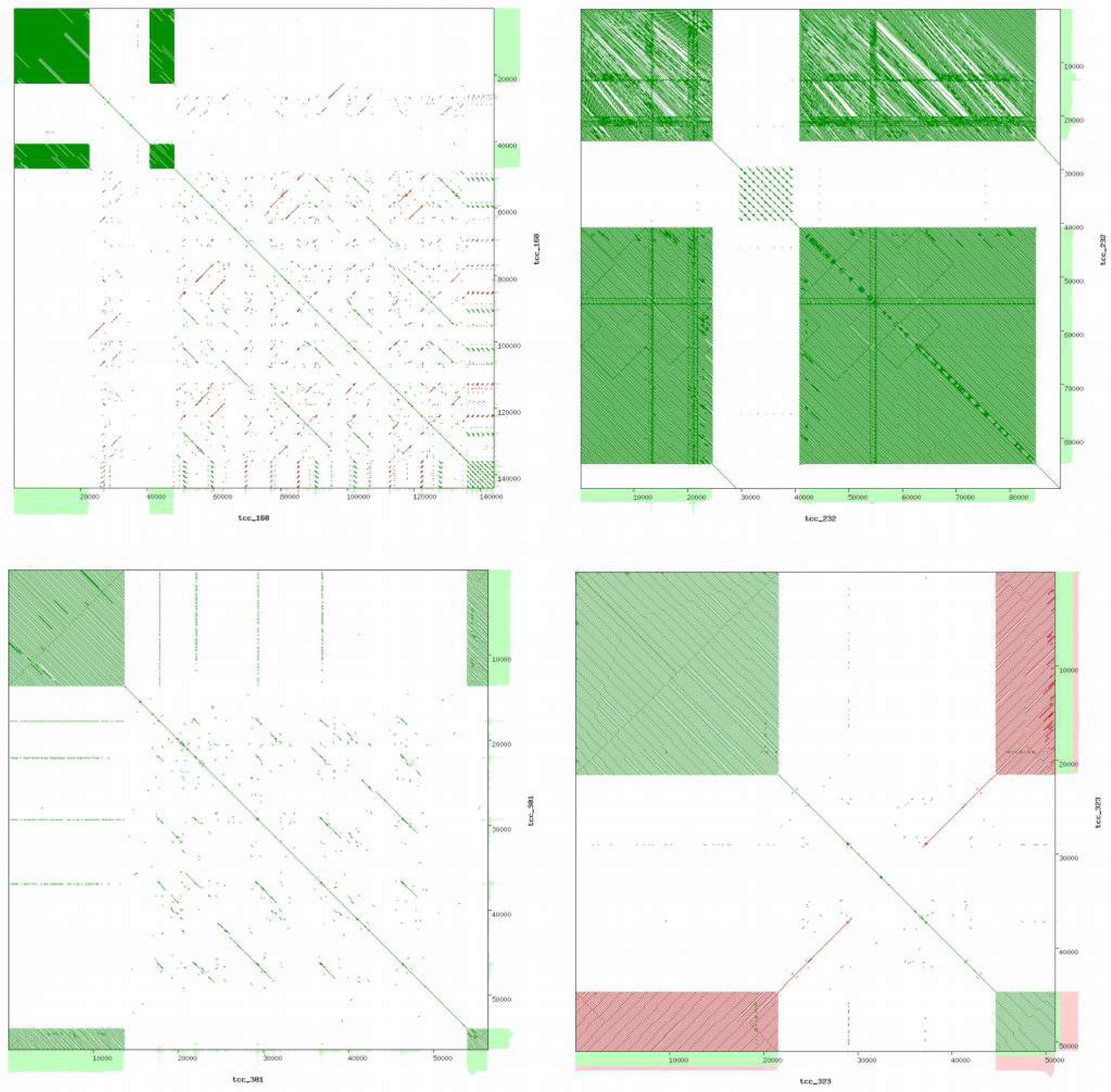

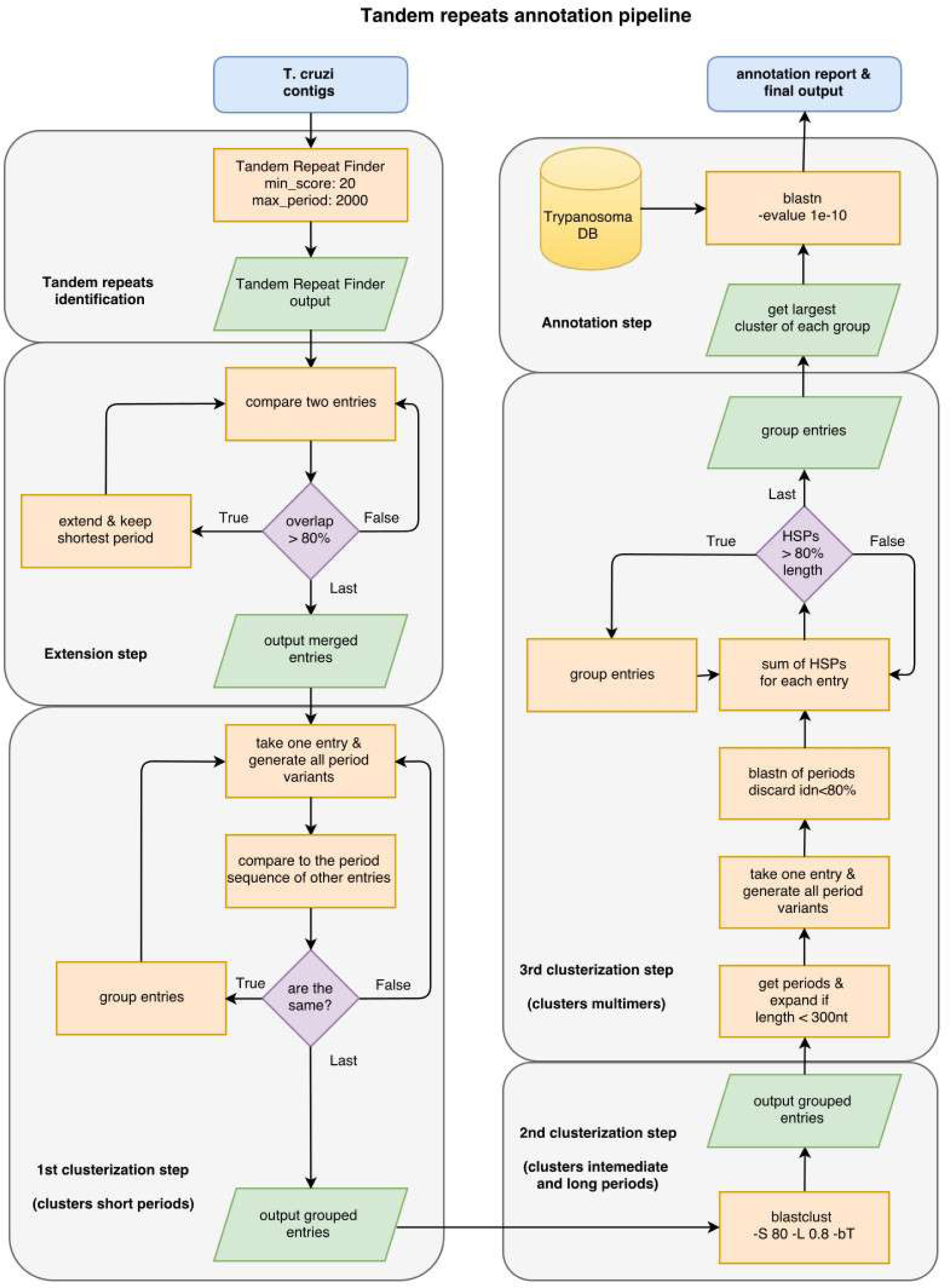

